# Investigating *Leptospira* dynamics in a multi-host community using an agent-based modeling approach

**DOI:** 10.1101/2022.06.27.497863

**Authors:** Aniruddha V. Belsare, Matthew E. Gompper, Meghan Mason, Claudia Munoz-Zanzi

## Abstract

Leptospirosis, a neglected bacterial zoonosis, is a global public health issue disproportionately affecting impoverished communities such as urban slums in the developing world. A variety of animal species, including peridomestic rodents and dogs, can be infected with different strains of leptospirosis. Humans contract leptospirosis via exposure to water or soil contaminated with the urine of infected animals. Due to the unavailability of safe and effective vaccines, preventive strategies mainly focus on minimizing human exposure to contaminated environment. In marginalized communities, this approach is ineffective due to infrastructure deficiencies and the difficulties in implementing sanitation and hygiene practices. Moreover, continuing expansion of urban slums worldwide will likely contribute to the increase in outbreaks of leptospirosis. Effective prevention of leptospirosis outbreaks will therefore require a thorough understanding of *Leptospira* transmission dynamics in impoverished, high-density settings. We developed the agent-based model MHMSLeptoDy to investigate *Leptospira* dynamics in a realistic, *in silico* high-density community of rodents, dogs, and human hosts, and two host-adapted *Leptospira* strains. Virtual explorations using MHMSLeptoDy were undertaken to evaluate alternate interventions and to assess the zoonotic transmission risk of leptospirosis. A key finding from model explorations is that rodents are the main contributors of rodent-adapted as well as dog-adapted strains in the environment, while dogs play an important role in distributing the rodent-adapted strain. Assessment of alternate leptospirosis control strategies using model iterations revealed that dog vaccination and rodent control interventions had the most impact on lowering human outbreak probability. Our open-source, customizable agent-based model provides a sophisticated mechanism to quantitatively evaluate nuanced intervention strategies and inform the design of rational, locally relevant leptospirosis control programs.

## INTRODUCTION

The complex ecology of multihost pathogens poses a major challenge for effective management of zoonotic diseases. In addition to the need to understand the role and importance of multiple host species, factors like pathogen genetic variability (multi-strain pathogens) or multiple modes of transmission (direct, environmental, vector-borne, etc.) can add to the complexity (1–3). Model-based approaches account for such complexities and facilitate investigation of pathogen transmission dynamics within a One Health framework by incorporating the interconnections between humans, animal hosts, and their environment.

Leptospirosis, a neglected zoonosis caused by bacteria in the genus *Leptospira*, is a case in point for model-based investigations. Humans are dead-end hosts for *Leptospira* and mainly contract the disease via exposure to water or soil contaminated with the urine of infected animals (4). Although rodents are recognized as primary reservoirs, a wide variety of species can also contribute to the cycling of leptospires in the environment, and some host species are believed to favor specific serovars or strains (5,6). Locale-specific host-strain associations and the environment in which they interact are therefore important considerations for designing and implementing effective leptospirosis control strategies.

In recent decades, leptospirosis outbreaks are increasingly reported from urban slum communities across the developing world (7–9). Limited infrastructure, poor sanitation, and inadequate waste management expose slum dwellers to environmental sources of contamination, thereby posing a higher risk of leptospirosis (8). Moreover, urban slums in developing countries support high densities of peridomestic and domestic species like rodents and dogs, that are known reservoirs or vectors of pathogenic *Leptospira* strains. Dogs in such settings are mostly free-roaming, with access to contaminated environments as well as a higher likelihood of contacting wild or peridomestic rodent species that may also harbor dog-adapted or other host species-adapted strains (10). Dogs can thus play a vital role in zoonotic transmission of leptospirosis, either as reservoir or maintenance hosts, or as vectors (11–14).

Globally, the public health impact of leptospirosis is estimated to be approximately one million cases per year resulting in >50,000 deaths (15). Current control measures have been nearly uniformly ineffective in addressing the public health threats posed by leptospirosis in developing countries (16). As safe and efficacious leptospirosis vaccines for humans are not available currently, preventive measures are limited and focus on minimizing human exposure to the environmental sources of contamination. In this context, education of proper sanitation and hygiene practices, as well as interventions that address infrastructure deficiencies are highly recommended but also difficult to implement in impoverished settings like urban slums. Thus, effective prevention of leptospirosis outbreaks in impoverished settings will require a more thorough understanding of the transmission dynamics in such communities, specifically the relative importance of dogs and rodents in zoonotic transmission of *Leptospira*. This understanding will be vital for identifying effective, locally relevant intervention strategies.

Agent-based (individual-based) models can simulate individual-level processes like host-host interactions, host-environment interactions, and the transmission of infectious agents as a stochastic by-product of such interactions. Moreover, such models can explicitly represent spatiotemporal elements of agent (host) interactions as well as agent - environment interactions and can thus provide valuable information about disease transmission dynamics. Here, we describe MHMSLeptoDy (Multi-host, Multi-serovar *Leptospira* Dynamics), an agent-based model for investigating the complex dynamics of humans, reservoir animal species, prevalent *Leptospira* strains and the environment in which they interact. We illustrate how MHMSLeptoDy can be used to assess the relative importance of various components that contribute to the zoonotic transmission risk of leptospirosis in urban slum settings. Insights gained from model explorations can be used to inform meaningful leptospirosis control strategies. MHMSLeptoDy has a user-friendly interface, is readily customizable (for instance, to non-urban settings or for other rodent pathogens of interest), and freely available for download from the CoMSES Net Computational Model Library, thereby facilitating its application for the design of defensible, locally relevant leptospirosis control strategies.

## MATERIALS AND METHODS

### Model description

The model, MHMSLeptoDy, was developed in NetLogo 6.1.1, a software platform for implementing agent-based models (17). Model code has been peer-reviewed and published as open access via website repository “Open ABM CoMSES Computational Model Library” (18). Detailed model description is provided following the Overview, Design concepts, Details (ODD) protocol for individual-based models in Supplementary Material. Model components and sequence of events are shown in Fig. S2.

The model landscape incorporates the general characteristics of an urban slum with high housing density and the presence of people, rodents and dogs, and is informed by data collected from an eco-epidemiological study of leptospirosis in Chile (19). Specifically, MHMSLeptoDy simulates population dynamics and behavior of rodents and dogs, and the emergent transmission of seroR (rodent adapted strain) and seroD (dog adapted strain).

Dogs in the model are essentially free-roaming, but some are closely associated with households (hence *‘owned’*) and potentially accessible for interventions. Dogs not associated with specific households are designated as *‘stray’* and are modeled to have increased susceptibility to infection (50% higher direct transmission probability) given an assumed poor health status compared to owned dogs.

The model also incorporates immune dynamics in infected and recovered hosts. Rodents infected with seroR remain persistently infected and continue to shed leptospires intermittently, whereas rodents recovering from seroD infection have lifelong immunity against seroD. Dogs recovering from seroD infection have immunity against seroD for a year, but dogs that recover from seroR infection have lifelong immunity against seroR. There is no cross-protection, so immunity against one strain does not protect the host from infection with the other strain. However, there is no co-infection simulated in the model, so a dog or rodent cannot simultaneously be infected with both strains.

Infected animals contaminate the model landscape with leptospires shed in their urine. Once in the environment, indirect transmission of the pathogen is possible to any host species, including humans. Intraspecific direct transmission also occurs for dogs and rodents. Daily movement of agents is not explicitly simulated in this model, but contamination of patches and the direct and indirect transmission probabilities are modeled to capture the movement pattern of agents in a typical day.

### Model evaluation and local sensitivity analysis

MHMSLeptoDy performance was evaluated by simulating the model without disease control intervention (baseline scenario). Human, rodent and dog populations were initialized during the model setup as described in the ODD protocol (Supplementary Materials). In brief, the model landscape was set to represent a slum community with 150 households over an area of ~ 0.3 km^2^. The human: dog ratio was set at 3 and the mean number of rodents per household was set at 9.5. The first year of the model run was without *Leptospira* infection in the model host community. In the first week of the second year, two percent of randomly selected stray dogs and two percent of randomly selected owned dogs were assigned seroD ‘shedder’ status and two percent each of stray and owned dogs were assigned ‘recovered’ from seroD infection status. Eight percent of randomly selected rats were assigned seroR ‘shedder’ status, and an additional two percent were designated latently infected with seroR status. After a one-year burn-out period, the model output file documented the following for a five-year period (year three to year seven): number of seroD human cases, number of seroR human cases, proportion of rodent contributed seroD human cases, proportion of dog contributed seroR human cases, number of seroD dog cases, number of seroR dog cases, number of seroR rodent cases, and number of seroD rodent cases. Average size of the rodent and dog populations during each year was also documented. To account for stochasticity in the model runs, model-generated data from 100 iterations was summarized for analysis.

The leptospirosis incidence rate per host week is calculated as (20):

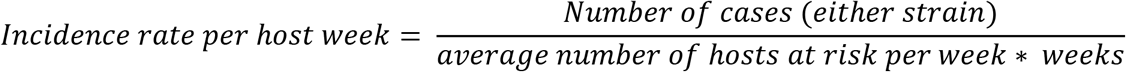

We performed a local sensitivity analysis of annual leptospirosis incidence in dogs and rodents over a period of 5 years. The model was set in the same manner as described above, and sensitivity values were generated for three key transmission parameters with most uncertainty, dog-to-dog transmission, rodent-to-rodent transmission, and environmental transmission, following Railsback and Grimm (21). A range was constructed for each parameter such that the lower (R^-^) and upper (R^+^) endpoints were ±15% of the reference value (R). For each parameter value, 100 iterations of MHMSLeptoDy were undertaken to generate sensitivity values. Sensitivities were calculated as

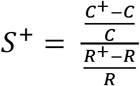

and

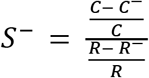

where C^-^, C and C^+^ are average annual incidence values (dogs or rodents) when a parameter is valued at R^-^, R and R^+^, respectively.

### Model Application: Evaluating Leptospirosis control interventions

MHMSLeptoDy was used to assess the impact of hypothetical interventions by modifying baselines conditions, such as reducing rodent or dog populations. For rodents, this intervention would mimic a rodent control program that was implemented with varying levels of sustained success (25% and 50% rodent population reduction compared to the baseline scenario). The interface slider *mean-num-rodents-perhh* was set at 7 for the 25% reduction scenario (RC25), and at 5 for the 50% reduction scenario (RC50).

For dogs, the intervention mimics a control program that targets general population reductions (stray and owned dogs) with varying levels of sustained success (25% and 50% reduction), which might occur via a variety of mechanisms, including culling, dog husbandry practices or policies that influence human ownership or acceptance of dogs. The interface slider *num-humans-per-dog* was set at 4 for the 25% reduction scenario (DC25), and at 6 for the 50% reduction scenario (DC50).

Additionally, dog vaccination was evaluated as a leptospirosis control intervention. Six dog vaccination scenarios were simulated with either monovalent vaccine providing protection against seroD only (MDV) or a bivalent vaccine (BDV) providing protection against both seroD and seroR. Three vaccination coverage scenarios were assessed: low at 25% coverage (MDV25/BDV25), medium at 50% (MDV50/BDV50) and high at 75% (MDV75/BDV75). We make a simplified assumption that vaccination results in full protective immunity against one or both the strains for one year and there is no cross-immunity for the MDV.

For each scenario, model-generated data from 100 iterations was summarized. The impact of each intervention was compared with the baseline scenario (no intervention, see Model Evaluation above) using incidence rate per host week calculated for dogs and rodents over a 5-year period. Persistence probability of a strain was calculated as the proportion of years with non-zero cases. Persistence probabilities were calculated for both strains in dog and rodent populations. Additionally, the public health impact of interventions was assessed by comparing the annual probability of human outbreak (probability of spillover into humans). For each strain, the annual probability of human outbreak was calculated as the proportion of years with non-zero human cases due to the strain. Annual overall probability of human outbreak is calculated as the proportion of years with non-zero human cases caused by either seroR or seroD.

## RESULTS

### Model evaluation and sensitivity analysis

Throughout the model run, MHMSLeptoDy landscape supported a mean weekly owned and stray dog population of 167 (± 20 SD) and a mean weekly population of 1437 (± 34 SD) rodents. The human population remained stable at ~750. After introduction in the second year, the rodent strain (seroR) persisted in the model host community in all iterations, while the dog strain (seroD) persisted in 98% iterations. The mean incidence rate of leptospirosis (seroR or seroD) in the rodent population was 0.044 per rodent-week. An average of 3103 (± 487 SD) seroR cases and an average of 206 (± 85 SD) seroD cases every documented in the model rodent population each year. The mean incidence rate of leptospirosis (seroR or seroD) in the dog population was 0.012 per dog-week. Interestingly, rodents contributed ~ 67% seroD leptospires in the environment while dogs contributed ~ 2% seroR leptospires in the environment.

An average of 52 (± 21 SD) seroD cases and 52 (± 13 SD) seroR cases were documented every year. The annual overall probability of human outbreak was 0.33, the annual probability of human outbreak for seroR was 0.23 and for seroD it was 0.17.

Local sensitivity analysis indicated that dog annual incidence was particularly sensitive to environmental transmission parameter, while rodent annual incidence was sensitive to rodent-to-rodent transmission parameter (Table 1).

**Table 1.** Local sensitivity analysis of annual leptospirosis incidence in dogs and rodents over a period of 5 years for three key transmission parameters, dog-to-dog transmission, rodent-to-rodent transmission, and environmental transmission.

| Parameter | Range | Rodent annual incidence |  | Dog annual incidence |  |
| --- | --- | --- | --- | --- | --- |
|  |  | S<sup>+</sup> | S<sup>-</sup> | S<sup>+</sup> | S<sup>-</sup> |
| dog-to-dog transmission | 0.0085, 0.01, 0.0115 | 0.09 | -0.03 | 0.33 | 0.01 |
| rodent-to-rodent transmission | 0.0085, 0.01, 0.0115 | 0.54 | 0.72 | 0.22 | 0.03 |
| environmental transmission | $1.8 \times 10^{-5}$, $2.12 \times 10^{-5}$, $2.44 \times 10^{-5}$ | 0.42 | 0.24 | 1.1 | 0.96 |

### Model application

#### Rodent control

For the 25% and 50% rodent population reduction scenarios (RC25 and RC50, respectively), the mean weekly population of rodents was 1059 ± 28 SD (~ 24% reduction compared to the baseline scenario) and 756 ± 21 SD (~47% reduction compared to the baseline scenario), respectively. Dog and human populations were similar in both rodent control and baseline scenarios.

SeroR persisted in both host taxa in all iterations for both rodent control scenarios, but seroD persistence was lower compared to the baseline scenario. Lower rodent populations resulted in lower incidence of leptospirosis in rodents as well as dogs (Table 2). The largest impact of rodent control interventions was on transmission of seroR in the rodent population as reflected in the lower number of infected rodents. Number of dogs infected with seroR, and number of rodents infected with seroD also decreased with a decrease in rodent populations, but no effect was evident on the transmission of seroD in dogs (Figure 1). Rodent contribution to environmental contamination with seroR and seroD also decreased with the intensity of rodent control (Table 3). Rodent control resulted in lower annual probability of human outbreak for both seroR and seroD, consequently, the overall annual probability of human outbreak was also lower for rodent control scenarios (Table 4).

**Table 2.** *Leptospira* strain dynamics in rodents and dogs for the model scenarios. For any scenario, persistence probability of a strain is calculated as the proportion of years with non-zero cases caused by the strain.

| Scenario | Rodents |  |  | Dogs |  |  |
| --- | --- | --- | --- | --- | --- | --- |
|  | SeroR persistence (%) | SeroD persistence (%) | Mean incidence (per rodent-week) | SeroR persistence (%) | SeroD persistence (%) | Mean incidence (per dog-week) |
| BL | 100 | 98 | 0.044 | 100 | 98 | 0.012 |
| RC25 | 100 | 93 | 0.031 | 100 | 93 | 0.011 |
| RC50 | 100 | 89 | 0.020 | 100 | 89 | 0.010 |
| DC25 | 100 | 86 | 0.042 | 100 | 84 | 0.010 |
| DC50 | 100 | 60 | 0.042 | 100 | 50 | 0.007 |
| MDV25 | 100 | 95 | 0.045 | 100 | 95 | 0.010 |
| BDV25 | 100 | 89 | 0.044 | 100 | 84 | 0.009 |
| MDV50 | 100 | 67 | 0.043 | 100 | 58 | 0.007 |
| BDV50 | 100 | 62 | 0.042 | 100 | 56 | 0.005 |
| MDV75 | 100 | 15 | 0.042 | 100 | 11 | 0.006 |
| BDV75 | 100 | 14 | 0.041 | 100 | 7 | 0.002 |

**Figure 1.**
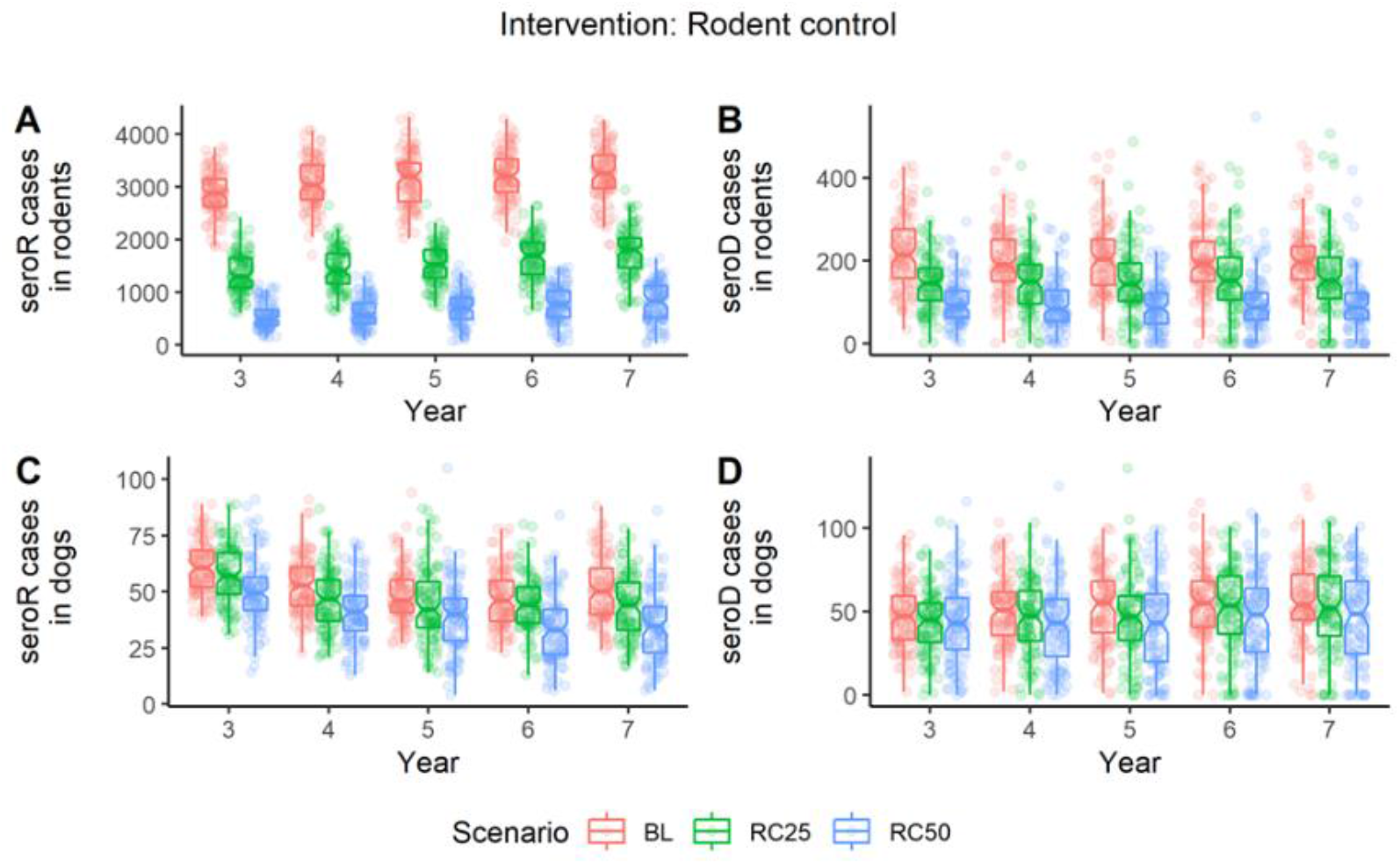
Impact of rodent control interventions on *Leptospira* serovar dynamics in MHMSLeptoDy landscape representing a marginalized urban setting. Number of infected hosts for each strain are compared between the baseline (BL; no intervention) scenario and the two rodent control scenarios (RC25 and RC50). Each circle represents the number of infected cases documented in a year for an iteration.

**Table 3.**
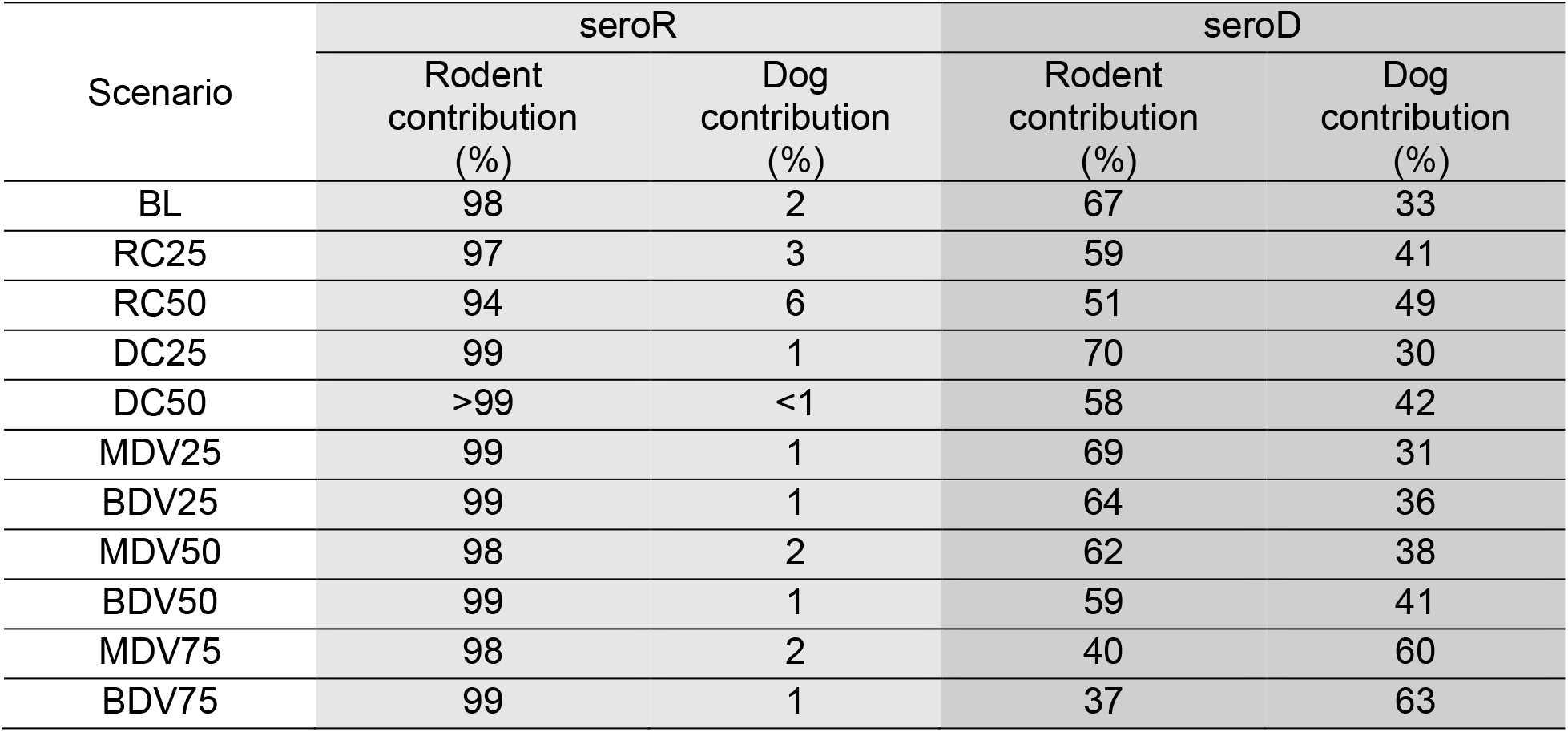
Strain-specific relative contribution of rodents and dogs to leptospires in the environment (pathogen pressure).

| Scenario | seroR |  | seroD |  |
| --- | --- | --- | --- | --- |
|  | Rodent contribution (%) | Dog contribution (%) | Rodent contribution (%) | Dog contribution (%) |
| BL | 98 | 2 | 67 | 33 |
| RC25 | 97 | 3 | 59 | 41 |
| RC50 | 94 | 6 | 51 | 49 |
| DC25 | 99 | 1 | 70 | 30 |
| DC50 | >99 | <1 | 58 | 42 |
| MDV25 | 99 | 1 | 69 | 31 |
| BDV25 | 99 | 1 | 64 | 36 |
| MDV50 | 98 | 2 | 62 | 38 |
| BDV50 | 99 | 1 | 59 | 41 |
| MDV75 | 98 | 2 | 40 | 60 |
| BDV75 | 99 | 1 | 37 | 63 |

**Table 4.** Strain-specific and overall leptospirosis outbreak probabilities in humans derived for model scenarios.

| Scenario | Human outbreak probability |  |  |
| --- | --- | --- | --- |
|  | seroR | seroD | Overall |
| BL | 0.23 | 0.17 | 0.33 |
| RC25 | 0.18 | 0.14 | 0.27 |
| RC50 | 0.11 | 0.12 | 0.20 |
| DC25 | 0.21 | 0.11 | 0.26 |
| DC50 | 0.20 | 0.05 | 0.23 |
| MDV25 | 0.19 | 0.14 | 0.27 |
| BDV25 | 0.20 | 0.13 | 0.27 |
| MDV50 | 0.21 | 0.09 | 0.26 |
| BDV50 | 0.18 | 0.07 | 0.23 |
| MDV75 | 0.21 | 0.04 | 0.23 |
| BDV75 | 0.16 | 0.02 | 0.18 |

#### Dog control

For the 25% and 50% dog population control scenarios (DC25 and DC50), the mean weekly population of dogs was 125 (± 18 SD) (~ 25% reduction in dog population compared to the baseline scenario) and 87 (± 14 SD) (~48% reduction compared to the baseline scenario), respectively. Rodent and human populations were similar in both dog control and baseline scenarios.

For both dog control scenarios, seroR persisted in both host taxa in all iterations, but seroD persistence was lower than the baseline scenario as well as the rodent control scenarios. Surprisingly, seroD persistence was higher in rodents compared to dogs for the dog control scenarios. Leptospirosis incidence in dogs decreased with increasing intensity of dog control, but in rodents, incidence of leptospirosis remained higher compared to the rodent control scenarios (Table 2). SeroR cases in rodents remained the same (Figure 2A) but seroR cases in dogs decreased for DC25 and DC50 scenarios (Figure 2B). SeroD cases in rodents and dogs also decreased with dog control, and a significant decline was documented for DC50 scenario (Figure 2B and 2D). Environmental contamination with seroR shed by dogs remained minimal (≤ 1%) in both dog control scenarios, and rodent shed seroD contamination was considerably lower for the DC50 scenario (Table 3).

**Figure 2.**
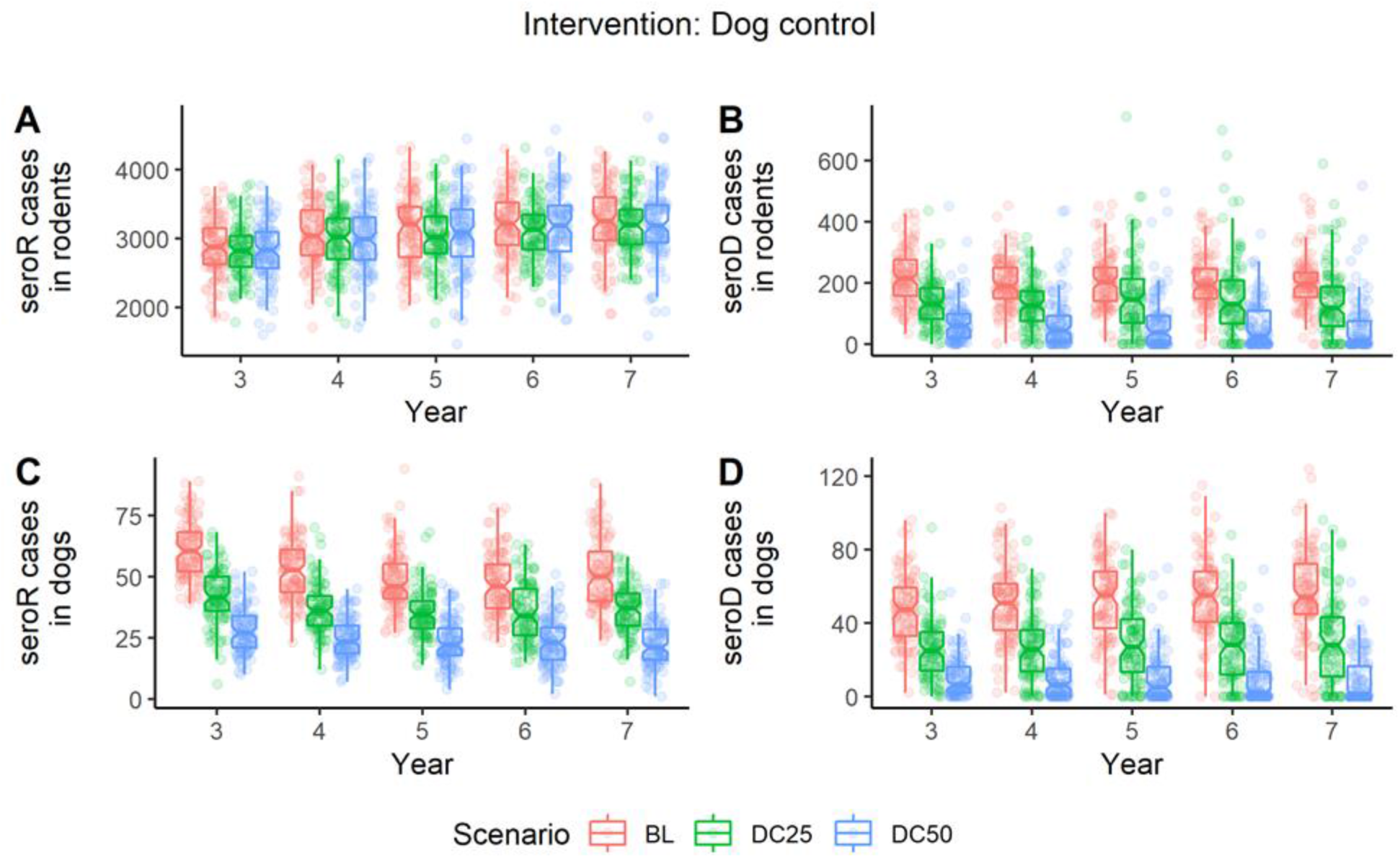
Impact of dog control interventions on *Leptospira* serovar dynamics in MHMSLeptoDy landscape representing a marginalized urban setting. Number of infected hosts for each strain are compared between the baseline (BL; no intervention) scenario and the two dog control scenarios (DC25 and DC50). Each circle represents the number of infected cases documented in a year for an iteration.

Dog control interventions lowered the annual probability of human outbreak for seroD, but the annual probability of human outbreak for seroR decreased only marginally. Dog control interventions lowered the overall annual probability of human outbreak of leptospirosis compared to the baseline scenario (Table 4).

#### Dog vaccination

For all dog vaccination scenarios, host populations in the model landscape were similar to the host populations in the baseline scenario. Dog vaccination did not affect seroR persistence in the model host community. In contrast, seroD persistence in dogs and rodents decreased with increasing vaccination coverage. Moreover, with the same coverage, vaccination with bivalent vaccine resulted in lower seroD persistence in both hosts compared to interventions with monovalent vaccines (Table 2). Leptospirosis incidence in dogs also followed the same pattern – increasing vaccination coverage resulted in decreasing leptospirosis incidence, and bivalent vaccines were more effective in lowering leptospirosis incidence compared to monovalent vaccines. However, leptospirosis incidence in rodents did not change notably, and remained above 0.040, similar to the baseline value of 0.044 (Table 2). Dog vaccination had no effect on seroR incidence in rodents but had a significant impact on seroD incidence in dogs and rodents, regardless of the vaccine type (Figure 3A, 3B, 3D). SeroR incidence in dogs decreased with increasing coverage of bivalent vaccine (Figure 3C). Environmental contamination with seroR shed by dogs remained low (≤ 2%) for all dog vaccination scenarios. Rodent shed seroD contamination decreased with increasing dog vaccination coverage and was lower for bivalent vaccine scenarios compared to the monovalent vaccine scenarios (Table 3).

**Figure 3.**
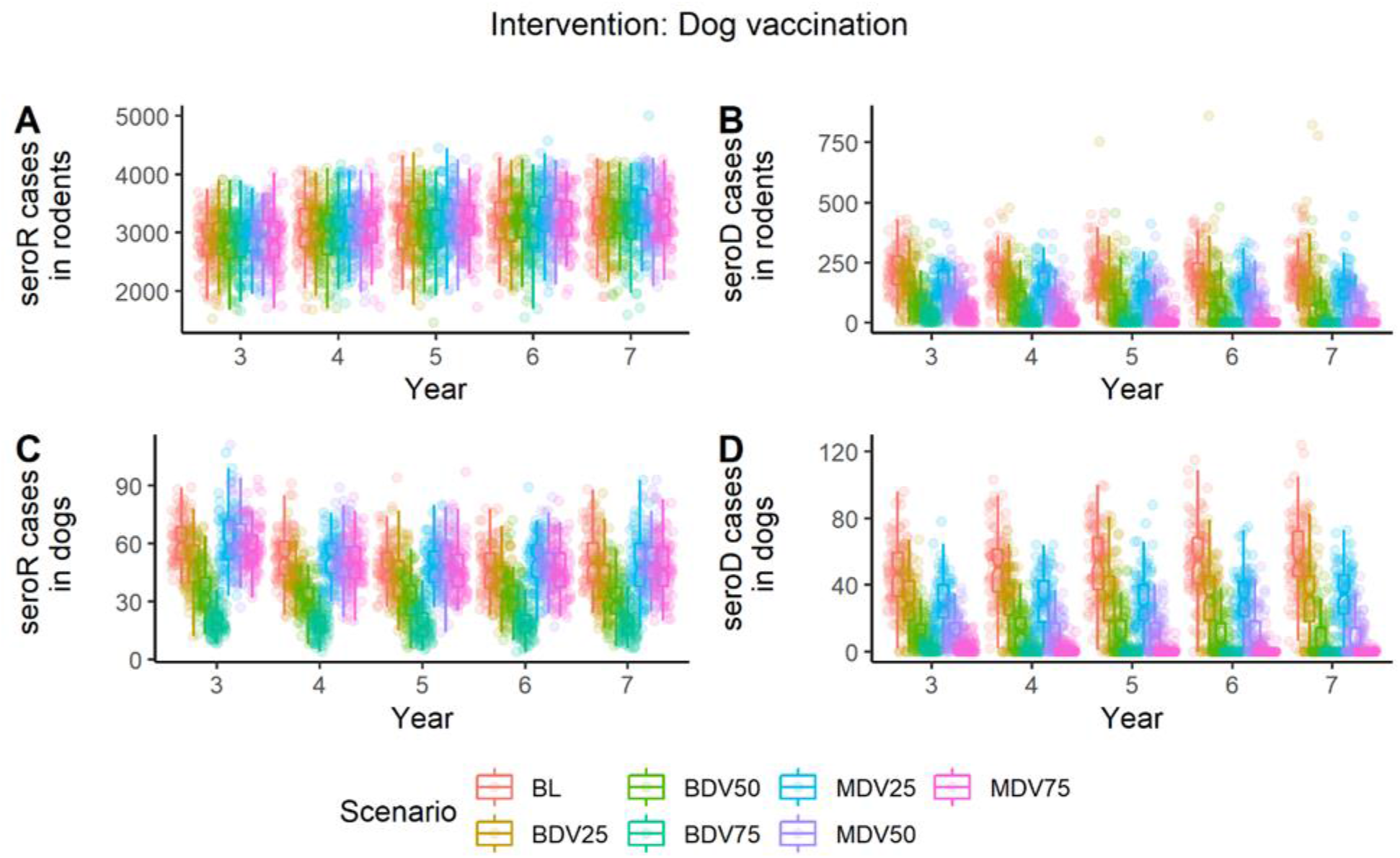
Impact of dog vaccination on *Leptospira* strain dynamics in MHMSLeptoDy landscape representing a marginalized urban setting. Number of infected hosts for each strain are compared between the baseline (BL; no intervention) scenario and the six dog vaccination scenarios. Each circle represents the number of infected cases documented in a year for an iteration.

The annual probability of seroD outbreak in humans decreased with increasing vaccine coverage, while only medium and high coverage bivalent vaccination scenarios lowered the annual probability of seroR outbreak in humans. The overall annual probability of human leptospirosis outbreak decreased with increasing vaccination coverage and was lower for bivalent vaccine scenarios compared to monovalent vaccine scenarios with same coverage (Table 4).

## DISCUSSION

Initial explorations using MHMSLeptoDy provide important insights into the dynamics of the complex leptospirosis system, specifically the relative importance of various components in the context of zoonotic transmission risk. It is generally recognized that most human leptospirosis cases are acquired through environmental contamination (22). Therefore, host contributions of leptospires to the environment at a given point in space and time (pathogen pressure) is an important indicator of the zoonotic transmission risk (23). A unique feature of MHMSLeptoDy is the ability to include and track transmission of different strains. Our assessments revealed rodents as the main contributors of leptospires in the environment for both strains. Rodents contributed > 90% of the seroR leptospires and > 50% of the seroD leptospires in the model landscape in all scenarios except the MDV75 scenario, where rodent contribution of seroD was 37%. Thus, researchers working with multiple strains might consider that while some strains may be ‘adapted’ to particular hosts, the importance of those hosts may be less than expected when viewed in the context of environmental contamination.

Across all scenarios, seroR persisted in both rodent and dog populations, although rodent control interventions resulted in lower incidence of seroR in both animal host populations. Without any intervention, persistence of seroD was high (98%) in the host community. Dog population control lowered seroD persistence in the population, as did dog vaccination. Intriguingly, seroD persistence in rodent population also decreased for all dog control and dog vaccination scenarios but was generally higher compared to seroD persistence in the dog population for the respective scenarios. For the low coverage dog vaccination with monovalent vaccine scenario (MDV25), seroD persisted in 95% of iterations in the dog population (31 ± 16) as well as the rodent population (147 ± 71). This highlights the limitations of interventions with low vaccination rates. Notably, such low dog vaccination coverage is commonly observed in both high and low-income settings (24,25). Independent of the coverage, rodents sustained circulation of seroD to some extent, thus playing an important role in the persistence of both strains in the host community. These findings are consistent with observations from published studies that document the important role played by rodents in the transmission of *Leptospira* in a multihost community (15).

The contribution of rodents in maintaining and transmitting the dog-adapted strain is mirrored in dogs contributing to the distribution of the rodent strain. Dogs played an important vectoring role for seroR, which contributed to the zoonotic transmission risk of leptospirosis in the model landscape. In the MDV75 scenario, vaccinated dogs were not protected against seroR, therefore a higher number of annual seroR cases were documented in the dog population (50 ± 13 SD) compared to BDV75 scenario (19 ± 8 SD), and this corresponded with the higher human outbreak probability (0.23) in the MDV75 scenario compared to the BDV75 scenario (0.18). SeroD cases in rodents and dogs, and seroR cases in rodents were comparable in both these scenarios.

Overall, the interventions that had the most impact on lowering human outbreak probability were dog vaccination and rodent control. These two interventions highlight mechanisms that can potentially reduce leptospirosis transmission risk in urban slum settings. The former approach focuses on reducing the incidence of leptospirosis in the dog population. The vaccine-induced herd immunity in the dog population possibly contributes to the decreased seroD spillover resulting in lower seroD incidence in rodents. Although the overall incidence of leptospirosis in rodents decreased minimally (from 0.044, ~ 3300 annual cases in the baseline scenario to 0.041, ~3047 annual cases), the human outbreak probability for this scenario was lower than all other scenarios.

The other effective approach focuses on lowering seroR incidence in rodents. Simulations with effective rodent control (sustained 50% reduction of rodents) resulted in a significant decrease in seroR incidence in the rodent population (682 ± 325 SD cases per annum compared to 3103 ± 487 SD cases per annum in the baseline scenario). SeroD incidence in rodents also decreased (97 ± 64 SD cases per annum compared to 206 ± 85 SD cases per annum in the baseline scenario), suggesting a density-dependent mechanism of *Leptospira* transmission in rodents in urban slum settings. The mean weekly incidence of leptospirosis in rodents (both strains) was 0.02, significantly lower than the mean weekly incidence of 0.044 for the baseline scenario. Rodent control resulted in ~50% lower transmission risk as seroR human outbreak probability was 0.11 compared to 0.23 for the baseline scenario, while the overall human outbreak probability was 0.20 compared to 0.33 for the baseline scenario.

An important caveat to the findings reported above is that interventions like rodent control, dog control and dog vaccination are generally difficult to implement or sustain in impoverished settings. Nonetheless, we contrasted these scenarios because insights gained from such model investigations can inform the design of more realistic and effective leptospirosis control strategies. Further, more nuanced scenarios can be readily evaluated using an adaptive management approach. For instance, model assessments can help identify a combination of sustainable, low intensity interventions (e.g., 10% rodent control, 10% dog control and 25% dog vaccination coverage) that effectively reduce the zoonotic transmission risk of leptospirosis in resource-limited settings.

Urban outbreaks of leptospirosis are likely to increase and intensify due to the continuing expansion of urban slums worldwide and the increasing effects of natural disasters and extreme weather events in such settings (8,26,27). Prevention of leptospirosis outbreaks will therefore remain a major public health priority in the foreseeable future. In resource-limited settings, an effective approach to improving disease control strategies for complex disease systems like leptospirosis is through the integration of formal models with management and related policy decision making. Such models, and agent-based models in particular, can facilitate integration of observational and experimental data, provide a sophisticated mechanism to explore the effects of uncertainty, and can be used to explore a range of intervention scenarios for informing rational disease control strategies. Moreover, virtual explorations using a formal disease transmission model presents an economical and time sensitive alternative to direct field investigations that are costly, require years to complete and are difficult to implement and sustain.

## Contributors

AVB contributed to conceptualization and development of the model code, output analysis and visualization, writing the original draft, reviewing and editing the final manuscript. MEG contributed to conceptualization of the model, investigation of model parameters, and reviewing and editing of the final manuscript. MM contributed to conceptualization of the model, writing of some components of the model code, investigation of parameters, partial writing of methods, and reviewing of final manuscript. CMZ contributed to conceptualization of the original idea and of the model framework, funding acquisition of the field data, investigation of parameters, and editing and reviewing of the final manuscript.

## Data sharing

Model code for MHMSLeptoDy is available for download via the CoMSES Net Computational Model Library: https://doi.org/10.25937/zp7k-bq40. Model output analyzed in this paper is available here: https://github.com/anyadoc/MHMSLeptoDy

## Acknowledgements

Initial work that produced the original idea on the modeling framework was supported by NSF EEID (Project 0913570). The Working Group in Mathematical modeling of *Leptospira* transmission and intervention strategies organized by CMZ and J. Velasco-Hernandez was supported by the National Institute for Mathematical and Biological Synthesis (NIMBioS). AVB was supported by NSF Award P20GM104420 (2017-2019) and the U.S. Fish and Wildlife Service through the Pittman-Robertson Wildlife Restoration Act Grant MI W-155-R (2019-2020). Computational analyses were facilitated by access to the New Mexico State University’s Discovery High Performance Computing Cluster.

