## Supplementary Material for "Investigating *Leptospira* dynamics in a multi-host community using an agent-based modeling approach"

SUPPLEMENTARY MATERIAL for Agents of change: Investigating *Leptospira* dynamics in a multi-host community using an agent-based modeling approach.

ODD for **MHMSLeptoDy**: Multi-host, multi-serovar *Leptospira* Dynamics Model

Aniruddha Belsare

Contributors: Meghan Mason, M.E. Gompper and C. Muñoz-Zanzi.

This model builds upon a *Leptospira* dynamics agent-based model developed by Meghan Mason as a component of her dissertation research.

**Background**

Leptospirosis is a neglected, bacterial zoonosis with worldwide distribution, but primarily a disease of poverty. This disease has a complex epidemiology. Rodents are thought to be the primary reservoir, but several pathogenic serovars (or strains) of bacteria in the genus *Leptospira* exist, and a variety of species may act as reservoirs for these strains. Transmission is mainly by direct contact of intact mucous membranes or abraded skin with urine, urine contaminated soil, water or urine-contaminated food (Sykes et al., 2011). Human infection is the result of direct or indirect contact with *Leptospira* in the urine of infected animal hosts, primarily livestock, dogs, and rodents (Levett, 2001).

There is increasing evidence that dogs and dog-adapted strain Canicola play an important role in the burden of leptospirosis in humans in marginalized urban communities. What is needed is a more thorough understanding of the transmission dynamics of *Leptospira* in these marginalized urban communities, specifically the relative importance of dogs and rodents in the transmission of *Leptospira* to humans. This understanding will be vital for identifying meaningful intervention strategies.

The theoretical community in MHMSLeptoDy is based on characteristics and data collected (number of hosts, households, and prevalence of leptospirosis) from the urban slum communities from an eco-epidemiological study of leptospirosis in Los Rios Region, Chile (Muñoz-Zanzi et al., 2014).

**Purpose**

The purpose of this agent-based model is to simulate the multi-host, multi-strain dynamics of leptospirosis. The overall goal is to support the design of meaningful interventions for the disease in marginalized, high-density community consisting of dogs, rodents, and humans.

**Entities, State Variables and Scales:**

*Entities:* MHMSLeptoDy has four entities: rodents, dogs, humans, and patches. Rodents, dogs, and humans are modeled as individuals occupying the patches. State variables for the 4 entities are described in Table 1.

**Table 1. State Variables and Scales**

| **Agent** | **State Variable** | **Meaning/Value** |
| --- | --- | --- |
| Patches | *hh-id*  *num-humans*  *num-odogs-here*  *dcc*  *c-acc*  *p-inf-rod*  *p-inf-dog*  *e-amount-seroR*  *e-amount-seroD*  *dc-seroR*  *rc-seroD* | ID number of the household on a patch (0 if not a household patch)  Number of humans belonging to a household on a patch  (0 if no household on the patch)  Maximum number of owned dogs on the patch  Dog carrying capacity of a patch  Current accommodation potential for dogs  Probability that a host on the patch will be infected  with rodent strain (range: 0-1)  Probability that a host on the patch will be infected  with dog strain (range: 0-1)  Count of leptospires of SeroR on the patch  Count of leptospires of SeroD on the patch  dog contributed seroR  rodent contributes seroD |
| Rodents | *r-start-patch*  *r-age*  *r-immune-status*  *r-days-incub-rod*  *r-days-shed-rod*  *r-days-incub-dog*  *r-days-shed-dog*  *r-days-recovered*  *r-trans-prob*  *d-trans-prob* | Center patch of the rodent’s home range  Rodent’s age (in weeks)  Rodent’s immune status (Susceptible/Infected/Shedding/Recovered)  for SeroR and SeroD  Counter: number of days the rodent is latently infected with SeroR  Counter: number of days the rodent is shedding SeroR  Counter: number of days the rodent is latently infected with SeroD  Counter: number of days the rodent is shedding SeroD  Counter: number of days in the recovered status  Probability of direct transmission of rodent strain (range: 0-1)  Probability of direct transmission of dog strain (range: 0-1) |
| Dogs | *status*  *start-patch*  *age*  *d-immune-status*  *d-days-incub-rod*  *d-days-shed-rod*  *d-days-incub-dog*  *d-days-shed-dog*  *d-days-recovered*  *r-trans-prob*  *d-trans-prob*  *d-death-prob*  *sus* | Whether owned (0) or stray (1)  Center patch of the dog’s home range  Dog’s age (in weeks)  Dog’s immune status (Susceptible/Infected/Shedding/Recovered)  for SeroR and SeroD  Counter: number of days the dog is latently infected with SeroR  Counter: number of days the dog is shedding SeroR  Counter: number of days the dog is latently infected with SeroD  Counter: number of days the dog is shedding SeroD  Counter: number of days in the recovered status  Probability of direct transmission of rodent strain (range: 0-1)  Probability of direct transmission of dog strain (range: 0-1)  Weekly probability of death for the dog (range: 0-1)  Probability adjustment for increased susceptibility of infection  for stray dogs (range: 0-1) |
| Humans | *h-immune-status*  *h-days-inf-rod*  *h-days-inf-dog* | Human agents’ immune status (Susceptible/Infected/Recovered)  for SeroR and SeroD  Counter: number of days the human has been infected with SeroR  Counter: number of days the human has been infected with SeroD |

*Spatial scale*: The model landscape represents one square kilometer area (143 x 143 patches, each patch equals 49 square meters), though the effective area depends on three user-specified parameters: ‘*num-households*’ (number of households), ‘*hh-area*’ (households and their surrounding patches) and ‘*community-area*’ (area of influence of households).

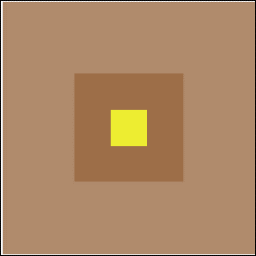

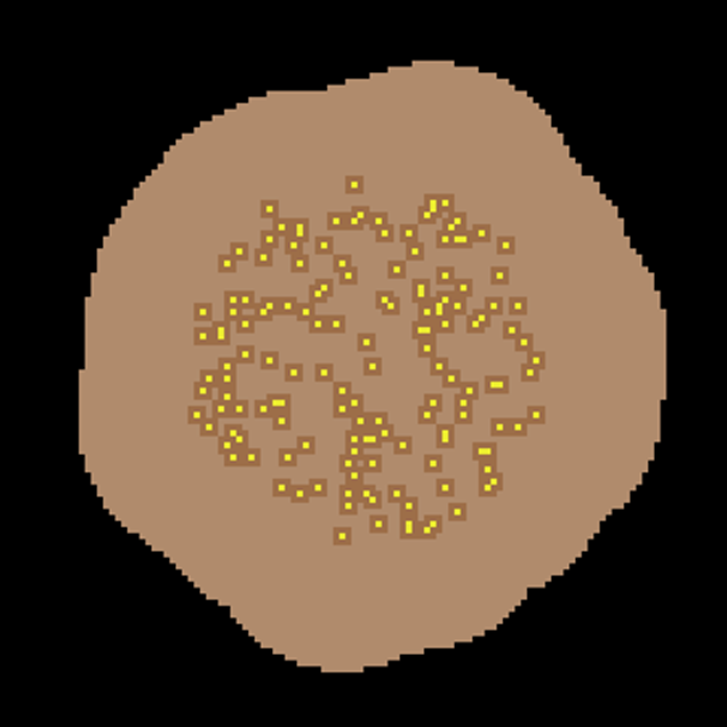

**Figure S1.** Model Landscape 'household' patches are designated as yellow patches. Eight patches around each household patch are designated as 'household-area' (dark brown patches). The effective model landscape for host-pathogen interactions includes the ‘*household*’, ‘*household-area*’ and the ‘*community area*’ (light brown patches + dark brown patches + yellow patches).

*Temporal scale*: MHMSLeptoDy has a weekly time step, and the model is simulated for 7 years. Only the host population dynamics are simulated in the 1^st^ year, and infection is introduced in the model host community in the 2^nd^ year. After a burnout period of 1 year (year 2), infection dynamics are recorded for years 3 through 7.

**Process overview and scheduling (Figure 2)**

During each time step (week):

1. Dogs and rodents execute three processes: ‘*die*’, ‘*birth*’ and ‘*age*’.
2. Humans execute ‘*die*’ and ‘*birth*’.
3. After 1 year (ticks > 51), *Leptospira* dynamics (‘*inf-dynamics*’) is executed 7 times every time step. Susceptible hosts have a probability of being directly infected with either seroD or seroR (*dog-to-dog* and *rodent-to-rodent*) or indirectly through the environment (contact with contaminated patches).
4. Infected rodents and dogs remain latently infected (not infectious) during the incubation period, followed by the infectious phase when leptospires are shed in urine (‘shedding’). Humans infected with a *Leptospira* strain are not infectious, and directly proceed to the recovered phase.
5. Rodents infected with rodent strain (seroR) remain persistently infected and continue to shed leptospires intermittently, whereas rodents recovering from dog strain (seroD) infection have lifelong immunity against seroD. Dogs recovering from dog strain (seroD) infection have immunity against seroD for a year, but dogs that recover from rodent strain (seroR) infection have lifelong immunity against seroR.
6. Leptospires present on the model patches decay. This process is also run 7 times every time-step.
7. Strain-specific prevalence in calculated for each host. Additionally, strain-specific incidence is calculated for humans.

**Figure S2: Process overview and scheduling.** Sequence of events during each step of the model are illustrated.

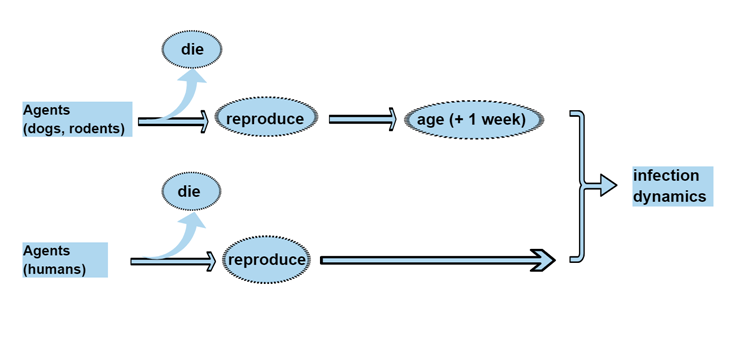

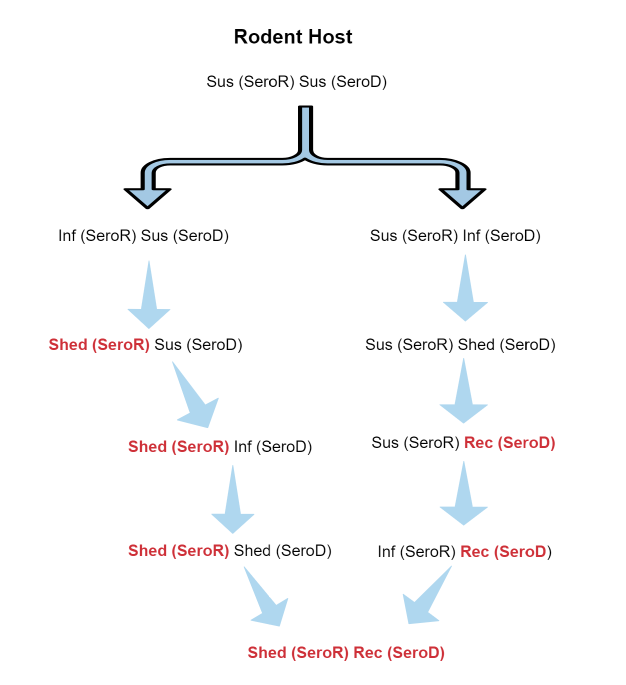

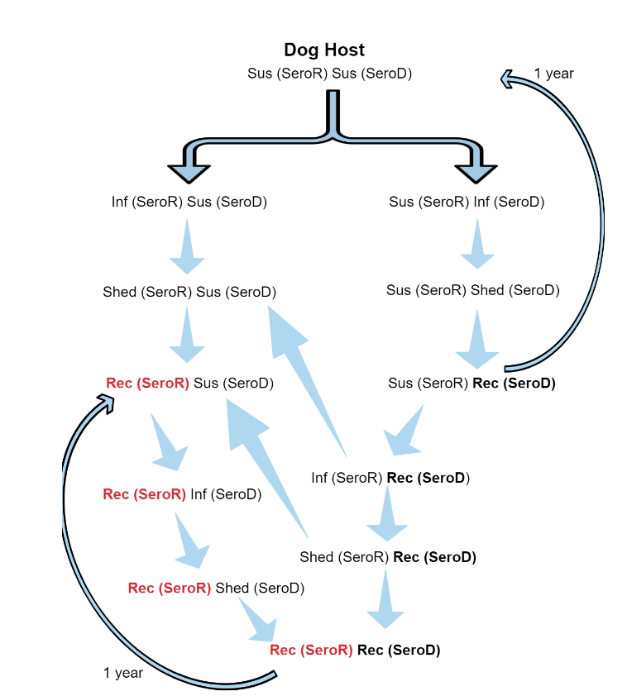

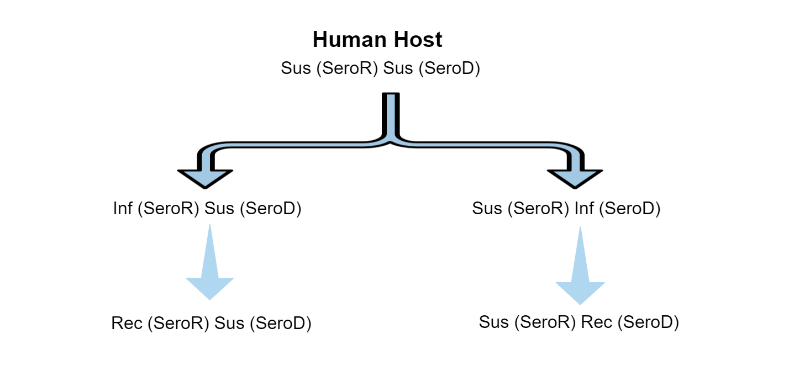

**Design Concepts**

*Emergence*: The age structure of rodent population emerges from this model. The population pattern of infection by each *Leptospira* strain in the three host species also emerges from the model.

*Adaptation*: The process of distribution of stray dogs in the model landscape is underpinned by the availability of vacant dog-supporting patches.

*Sensing*: Sub-adult stray dogs find their home-patch by ‘sensing’ vacant patch clusters in the vicinity of a patch.

*Interaction*: Host-strain interactions are underpinned by host immune status.

*Stochasticity*: Host distribution in the model landscape, initial age distribution of dog and rodent hosts, initial assignment of infected or shedding status are modeled as stochastic processes. The mortality and birth probabilities for dog and rodent hosts are derived from mortality and birth rates respectively, but the process of selection of individuals to implement mortality or birth is stochastic. The order in which hosts are exposed to infectious hosts and leptospires in the environmental is also stochastic.

*Observation*: Abundance and distribution of hosts (rodents, stray dogs and owned dogs) is updated and displayed (graphs) as the model executes. Prevalence of seroR and seroD in rodents, stray dogs and owned dogs is displayed graphically. Additionally, graphs of leptospirosis incidence and prevalence (both strains) in humans are also displayed on the model interface. Monitors display human incidence per 10,000 for both seroR and seroD. Additionally, the following model outputs are also recorded in a file titled ‘HostLeptoIncidence.csv’: i) seroD annual incidence per 10000 humans, ii) seroR annual incidence per 10000 humans, iii) annual number of cases due to seroD in dogs, iv) annual number of cases due to seroR in dogs, v) annual number of cases due to seroR in rodents, and vi) annual number of cases due to seroD in rodents.

**Initialization**

***Households and humans***: The user specifies the number of households desired, the ‘*hh-area*’ (households and their surrounding patches) and ‘*community-area*’ (area of influence of households). All patches are clear of contamination with leptospires.

Human agents are associated with household patches. Each household patch is initiated with a mean of 4 humans (range 1 - 11) (Mason et al., 2015). Distribution of human agents is maintained within this range throughout the model simulation.

***Dogs***: The carrying capacity of dogs is set at 2500 per square kilometer (extrapolated from Acosta-Jamett, 2010) (Acosta-Jamett et al., 2010). The user defines the human: dog ratio (*num-humans-per-dog*) in the community. Dogs are essentially free-roaming, but some are closely associated with households (hence ‘*owned’*) and potentially accessible for interventions. Dogs not associated with households are designated as ‘*stray’* dogs - these dogs are still dependent on the households (food, garbage, shelter) and so remain in the ‘*community-area*’. Owned: stray ratio is set at 1:1 (Meghan Mason, personal observation, November 2013).

Owned dogs have a household patch (yellow) as their '*home-patch*', while stray dogs can have any non-household patch (dark brown and light brown) as their '*home-patch*'. Owned dogs are distributed based on household dog ownership data from Chile: 47% of households do not own dogs, 27% own one dog, 11% own two dogs, 8% own three dogs, 5% own four dogs, and 2% own five dogs (Acosta-Jamett et al., 2010).

***Rodents***: The user defines the number of rodents per household in the community and the rodents are randomly distributed across ‘*hh-area*’ patches in the community area.

Transmission parameters: The user defines the probability of direct transmission from *rodent-to-rodent* and from *dog-to-dog*. The proportion of leptospires surviving each day is also user-defined (*decay-factor*).

The model is run for the first year with no infection. Infection is introduced in the first week of second year of the model run. Two percent of stray dogs and 2 % of owned dogs change their immune status to shedders (of seroD) and 2% each of stray and owned dogs are in the recovered stage (from seroD). Eight percent of rats change their status to seroR shedders and 2% are latently infected with seroR.

**Submodels**

1) **od-birth** / **sd-birth** / **od-die** / **sd-die**

We assume a male-skewed sex ratio of 63% in the adult dog population, so 37% of dogs were deemed females (Gompper, 2015). Fifty percent of adult females reproduce per year with an average litter size of 5, or 2.5 puppies per adult female per year. Birth occurs over a 16-week period between September and December annually. Translated into a weekly probability (1 - exp(-2.5/16)), the weekly birth probability for an adult female dog during the breeding season is 0.1447. The baseline weekly birth rate is adjusted by the proportion of initial number of dogs in a category (owned/stray) compared to their current number.

*p(birth) = 0.1447 ** $\frac{Initial number of owned or stray dogs}{Current number of owned or stray dogs}$

All newborn pups are in the susceptible class for both the strains (*d-immune-status* “ss”). The start-patch is the same as the mothers.

The weekly mortality probability for dogs is derived assuming an annual dog population turnover of 30%. We first converted the annual rate into instantaneous rate:

Instantaneous mortality rate = ln ( 1 – 0.3) = 0.3567

The instantaneous mortality rate was then converted into a weekly mortality probability using the equation:

probability (p) = 1 – exp(-rt)	(Briggs et al., 2006)

where r = instantaneous rate and t = period of interest.

The weekly mortality probability derived using the above equation is 0.00684, and this probability is same for both stray and owned dogs. To simulate density dependence, the baseline probability of death is adjusted by the proportion of current number of dogs in a category (owned/stray) compared to their initial number.

*p(death) = 0.00684 ** $\frac{Current number of owned or stray dogs}{Initial number of owned or stray dogs}$

Dogs infected with the rodent strain (seroR) have an increased probability of death (10% of the baseline probability). Patches subsidizing the dog become available (*change-av-dogs*) and the patch variable *c-acc* (current accommodation potential for dogs) is updated accordingly.

2) **r-die** / **r-birth**

Rodents are considered adults after two months, and half of the newborn rodents do not survive to adulthood. The weekly probability of death for juvenile rodents derived using the method described above is 0.0830. The weekly probability of death for adult rodents is 0.0957, derived from 80% of the rodents who survived beyond two months (Glass et al., 1989). The death probabilities are density dependent, adjusted by the proportion of current number of rodents compared to the initial number.

*p(death – juvenile) = 0.0830 ** $\frac{Current number of rodents}{Initial number of rodents}$

*p(death – adult) = 0.0957 ** $\frac{Current number of rodents}{Initial number of rodents}$

Rodents reproduce year-round, and an estimated 12-48 offspring are born per year per female rodent. The weekly probability of reproduction is set to a higher value for 26 weeks and to a lower value for the remaining year.

48 offspring per rodent female per year = 0.6027 weekly birth probability

12 offspring per rodent female per year = 0.2061 weekly birth probability

Twenty five percent of the total rodent population is modeled to reproduce at any time to reflect the age and sex structure of the population (A. Previtali, personal communication). Density dependence is factored in by scaling the weekly probability with the proportion of initial number of rodents to the current number of rodents.

*p(birth - high) = 0.6027 ** $\frac{Initial number of rodents}{Current number of rodents}$

*p(birth - low) = 0.2061 ** $\frac{Initial number of rodents}{Current number of rodents}$

3) **h-die** / **h-birth**

The model assumes that there is no migration into or out of the marginalized urban community. The human crude birth rate per 1,000 in Chile is 13.9 (World Health Organization, 2016), which translates to a weekly reproductive probability of 0.000266 (see method described under od-birth/sd-birth/od-die/sd-die submodel). There is no assumed seasonality to human births, so the weekly probability of death is also 0.000266. Human probability of death also increases by 10% when infected with *Leptospira* strains. As with the other agents, the more humans there are in the population, the more frequently deaths occur, and the less frequently births occur:

*p(birth) = 0.000266 ** $\frac{Initial number of humans}{Current number of humans}$

*p(death) = 0.000266 ** $\frac{Current number of humans}{Initial number of humans}$

4) **find-dohh**

At 6 months of age, pups born to owned dogs seek a dog-owning household that has a vacancy. A quarter of all the dog-owning households are scanned by the dog. If successful, the dog remains in the owned category and moves to the household patch with a vacancy. Otherwise, it changes the status to stray dog.

5) **find-sdog-hp**

Stray dogs find their home-patch by randomly scanning 10% of all household area patches for vacancy (enough dog subsidizing patches in the vicinity). If such a patch is found, the dog moves to this patch. If unsuccessful, the dog will do a random scan of all patches and move to the first vacant patch.

6) **chk-av-dogs**

A patch executing this submodel checks for available dog subsidizing patches in the vicinity (radius of 6 patches, ~ 42 meters), and if at least 9 such patches are found, the patch updates its capacity to accommodate a new dog (*c-acc*). The number of dog subsidizing patches required to support a dog is determined by dividing the slum area (total dog subsidizing patches in the model landscape) by the dog carrying capacity (~2500 per km^2^).

7) **change-av-dogs**

When a patch ‘acquires’ a dog, this submodel is executed to change the availability of 9 patches in the vicinity.

8) **inf-dynamics**

Dogs, rodents and humans execute the infection dynamics submodels (*d-get-infected*, *r-get-infected*, *h-get-infected* respectively) seven times every time step. Figure 2 illustrates the immune-status transition for the three hosts. Patches execute the e-decay submodel seven times every time step.

***d-get-infected***

Dogs susceptible to a strain can get infected directly from another dog (dog to dog transmission) or indirectly from a contaminated patch. Stray dogs are modeled to have increased susceptibility (50% higher direct transmission probability). Transmission probability is calculated as the product of total number of potential contacts with infectious dogs per day and the dog-to-dog transmission probability.

Potential number of infectious dogs contacted = infectious dogs on home patch + random number of infectious dogs on neighboring patches + random number of infectious dogs on a cluster of patches within a 20-patch radius (~140 meters). Similarly, environmental transmission (from contaminated patches) is modeled as the sum of probabilities of indirect transmission for the home patch, for randomly selected neighboring patches (0 to 8), and for a randomly selected cluster of up to 9 patches in a 20-patch radius. This process attempts to capture the movement pattern of dogs in their home range during a typical day.

During each iteration of the submodel *d-get-infected*, the sequence of exposure to the two serotypes (direct transmission by seroR infectious dogs, direct transmission by SeroD infectious dogs, indirect transmission seroR and indirect transmission SeroD) is randomly shuffled.

Dogs are latently infected for a duration of 3-21 days (seroR) or 3-14 days (seroD) (Ahmed et al., 2012; Greenlee et al., 2005; Sykes et al., 2011). After the latent infection period, the dog begins to shed leptospires into the environment. Forty percent of the total leptospires are shed on the home-patch, 30 % are shed on one or more of the neighboring patches and 30% are shed on a cluster of patches (1 – 8 patches) within a 20-patch radius (*dog-shed-dogsero* / *dog-shed-rodsero*).

The peak amount of leptospires to be shed by the dog is selected from a distribution constructed from the lab-based estimate of 5.9 x 10^4^ leptospires shed per mL (Rojas et al., 2010). To translate the concentration into a quantity for the model, the amount of urine shed per dog per day, estimated at 60mL per kg body weight was used (Campbell & Chapman, 2000). This was combined with the average weight of dogs observed on-site in urban slums of 15kg (Mason, personal observation, 2013), to obtain an average of 900mL of urine shed per dog per day in the model. The peak quantity of leptospires shed by dogs will be randomly selected from a normal distribution (mean 5.9 x 10^4^, sd 10^4^) x 900 mL per day. It takes three days after the incubation period limit has been reached for a dog to begin shedding the peak quantity of leptospires into the environment.

The duration of shedding will be longer for SeroD relative to SeroR, modeled as a more gradual decay function for SeroD relative to SeroR. The shedding functions based on the peak amount selected from the normal distribution (*peak*) for the amount of leptospires shed per day (A) for a dog (*d*) are shown below where *t* indicates the number of days the dog has been shedding and *c* is the decay rate. The decay rate is randomly selected from 0.15 to 0.35 for seroD and 0.3 to 0.7 for seroR. After the incubation period, a sharp rise in the quantity of leptospires in urine is noted (Bharti et al., 2003; Haake & Levett, 2015; Ko et al., 2009; Rojas et al., 2010), represented in this model by a linear progression from 0 leptospires to the peak quantity of leptospires shed, over the course of three days.

if *t* = 0,1,2,3: $A_{SeroD}^{d}$ = $A_{SeroD(peak)}^{d}$ * $(\frac{t}{3}$) and $A_{SeroR}^{d}$ = $A_{SeroR(peak)}^{d}$ * $(\frac{t}{3}$)

if *t* > 3: $A_{SeroD}^{d}$ = $A_{SeroD(peak)}^{d}$ * $e^{(\left( t-3 \right)*(-c))}$ and $A_{SeroR}^{d}$ = $A_{SeroR(peak)}^{d}$ * $e^{(\left( t-3 \right)*(-c))}$

When the quantity of leptospires shed drops below 100 leptospires per day, a dog is considered recovered and immune from the strain with which it was infected. Immunity is lifelong against seroR for dogs (Evangelista & Coburn, 2011; World Health Organization & International Leptospirosis Society, 2003), and immunity lasts one-year against seroD, as extrapolated from vaccine studies that suggest one-year protection with artificial immunity (Goldstein, 2010; Klaasen et al., 2014; Minke et al., 2009; Schreiber et al., 2005).

***r-get-infected***

Rodents susceptible to a strain can get infected directly from another rodent (rodent to rodent transmission) or indirectly from a contaminated patch. Transmission probability is calculated as total number of potential contacts with infectious rodents (for a strain) per day times the rodent-to-rodent transmission probability.

Potential number of infectious rodents contacted = infectious rodents on home patch + random number of infectious rodents on neighboring patches + random number of infectious rodents on a cluster of patches within a radius of the rodent’s home-range. Rodent home-range is set to have a radius of up to 1.5 patches (~10-meter radius). This process attempts to simulate the rodent contact pattern during a typical day.

Transmission (from contaminated patches) is modeled as the sum of probabilities of indirect transmission for the home patch, for randomly selected neighboring patches (0 to 8), and for a randomly selected cluster of up to 9 patches in the rodent’s home range.

During each iteration of the submodel *r-get-infected*, the sequence of exposure to the two serotypes (direct transmission by seroR infectious rodents, direct transmission by seroD infectious rodents, indirect transmission seroR and indirect transmission seroD) is randomly shuffled.

Rodents are latently infected for a duration of 7-14 days for seroR and seroD (Athanazio et al., 2008; Rojas et al., 2010). After the latent infection period, the rodent begins to shed leptospires into the environment. Seventy percent of the total leptospires are shed on the home-patch and 30 % are shed on one or more of the neighboring patches (1 – 8 patches) (*rod-shed-dogsero* / *rod-shed-rodsero*).

The peak amount of leptospires shed per mL of urine by rodents is randomly selected from a normal random distribution (mean 6.1 x 10^6^ leptospires, standard deviation 1.5 x 10^6^) based on shedding by Norway rats in Brazilian slum communities (Costa et al., 2015). In the eco-epidemiology of leptospirosis study, the average weight of rodents in marginalized urban communities was 138.4g (Munoz-Zanzi, unpublished data). If a rodent excretes 5.5 mL of urine per 100g body weight per day (Pass & Freeth, 1993), then the rodents in this model shed 7.59 mL of urine per day on average, yielding a randomly selected peak value from a normal distribution (mean 6.1 x 10^6^, sd 1.5 x 10^6^) * 7.59 mL. It takes three days after the incubation period limit has been reached for a rodent to begin shedding the peak quantity of leptospires into the environment.

Long-term shedding of leptospires by rodents is commonly reported (Monahan et al., 2008; Plank & Dean, 2000; Rojas et al., 2010), and therefore the shedding functions will be different when a rodent is infected with the rodent-adapted servoar as compared to the dog-adapted strain. The shedding functions based on the peak amount selected from the normal distribution (*peak*) for the amount of leptospires shed per day (A) for a rodent (*r*) are shown below where *t* indicates the number of days the dog has been shedding that strain and *c* is the decay rate. The decay rate is randomly selected 0.3 to 0.7 for SeroD and 0.15 to 0.35 for SeroR, allowing for a longer duration of shedding for the rodent-adapted strain. As with the dogs, a sharp increase in the quantity of leptospires shed occurs over the first three days of shedding:

if *t* = 0,1,2,3: $A_{SeroR}^{r}$ = $A_{SeroR(peak)}^{r}$ * $(\frac{t}{3}$) and $A_{SeroD}^{r}$ = $A_{SeroD(peak)}^{r}$ * $(\frac{t}{3}$)

if *t* > 3: $A_{SeroR}^{r}$ = $A_{SeroR(peak)}^{r}$ * $e^{(\left( t-3 \right)*(-c))}$ and $A_{SeroD}^{r}$ = $A_{SeroD(peak)}^{r}$ * $e^{(\left( t-3 \right)*(-c))}$

Rodents infected with and shedding SeroD are considered recovered when the quantity of leptospires shed drops below 100 leptospires per day, and the rodent has a lifelong immunity against SeroD. Rodents infected with SeroR remain persistently infected and shed leptospires intermittently. Immunity is lifelong against seroD, but a rodent is immediately susceptible again to infection seroR.

***h-get-infected***

Direct transmission to humans is not modeled, and so all infections are assumed to occur as a result of contact with a contaminated environment. To reflect protective measures that humans may take to protect themselves from zoonotic disease transmission from the environment, the probability of infection is reduced by a factor of 100 (Pass & Freeth, 1993). The indirect transmission probability for each strain is a sum of patch transmission probabilities of the home patch, of up to 9 random patches in a radius of ~ 10 meters and of up to 9 random patches in a radius of 350 meters.

p(infected)_SeroR_ = 0.01/10^9^ * *number of leptospires* _SeroR_

p(infected)_SeroD_ = 0.01/10^9^ * *number of leptospires* _SeroD_

Once infected, it is unlikely the human contributes to additional incidence of the disease in the community – humans are considered dead-end hosts for leptospirosis, even though they may shed (Ganoza et al., 2010; Ko et al., 2009). A human will recover after one month of being infected, and immunity to the infecting strain will be lifelong.

***e-decay***

Infectious rodents and dogs (Ganoza et al., 2010)(Ganoza et al., 2010)(Ganoza et al., 2010)(Ganoza et al., 2010)excrete leptospires into the environment in the amount indicated by the shedding equations (Ganoza et al., 2010). The leptospires that they shed are added to the number of leptospires already on the patch. Only a portion of the live leptospires survive at each time-step. A previous model for leptospirosis transmission in the rodent population used varying rates of decay in the environment of 5% to 20% per day (Holt et al., 2006). Therefore, each day, 5% of the current number of leptospires on a given patch will die until fewer than 1000 leptospires exist, and a patch will no longer be considered contaminated and contact with that patch would not be sufficient to produce infection in a host.
